# Direct measurement of PET hydrolase interfacial kinetics reveals catalysis-independent surface remodelling

**DOI:** 10.64898/2026.09.02.749009

**Authors:** Kailey J. Petz, Arnaud Boudigou, Laura E. Dickson, Benoît H. Lessard, Adam M. Damry

## Abstract

How PET hydrolases engage solid plastic has been inferred almost entirely from soluble analogs and surfactant-stabilised nanoparticle suspensions, and reported affinities span orders of magnitude. Here we measure it directly, depositing thin amorphous PET films onto gold surface plasmon resonance chips and following enzyme binding to authentic polymer in real time. Four PET hydrolases, LCC, LCC-ICCG, *Is*PETase-EHA and *Tf*Cut2, all bind with nanomolar apparent affinity and surface residence half-lives of tens of minutes. LCC-ICCG dissociates at 4.0 × 10⁻⁴ s⁻¹, within twofold of its *k*_cat_ measured on authentic PET, indicating that turnover is limited by disengagement rather than by ester hydrolysis. During association, we observe non-monophasic responses for these enzymes, including, for several, a signal fall while enzyme is still flowing over the surface. This decline persists in a catalytically inactivated variant, is absent in control proteins and a structurally homologous non-plastic degrading cutinase, and cannot arise from movement of the bound enzyme alone. The polymer surface must therefore be altered non-catalytically by PET hydrolase binding. The process runs at different rates for different enzymes on an identical film, is suppressed by dilute Triton X-100, and is modulated biphasically by PET degradation products. Altogether, we conclude that enzymatic PET depolymerisation involves a catalysis-independent chain-mobilisation step that soluble assays cannot report.

## Introduction

Plastics are widely used in modern society, but due to their long lifespan in the environment, they are remarkably persistent pollutants. Efforts to curtail plastic waste have turned to plastic degrading enzymes (PDEs) to combat plastic pollution and develop improved industrial recycling processes. The most efficient known PDEs are primarily active on PET and include variants of natural PDEs such as *Is*PETase, LCC cutinase, and related type I cutinases (henceforth collectively referred to as PET hydrolases).^1–3^ These enzymes are unusually efficient at degrading PET despite a low overall catalytic efficiency relative to other hydrolases.^4,5^ Due to a lack of ultrahigh-throughput screening assays for PDE activity, most engineering efforts seeking to improve these enzymes’ activity have focused on small mutant libraries. However, with a relative lack of mechanistic understanding of molecular interactions between PDEs and plastics, these engineering efforts have proven difficult. Indeed, while some studies report PET hydrolase activity enhancements of one to two orders of magnitude, as seen for example with the engineered enzymes FAST-PETase, LCC-ICCG, and HotPETase,^1,6,7^ these are often the result of increased thermostability rather than catalytic efficiency. Further convoluting engineering efforts, these enzymes possess highly epistatic mutational landscapes ^8,9^ and many PET hydrolase engineering efforts fail to yield substantially improved variants, reflecting unresolved questions about the molecular basis of PET recognition and turnover.^5,10,11^

As plastics are generally difficult to handle in aqueous systems, experimental insight into PET-PET hydrolase interactions remains limited. Many catalytic PET hydrolase studies and engineering campaigns rely on small molecule analogs like *p*-nitrophenyl esters and PET trimers (henceforth “3PET”) to simplify detection. However, these analogs poorly mimic PET’s structure, failing to capture its surface characteristics.^12–14^ When authentic PET is used, PET hydrolase kinetics are often described indirectly, with catalytic rates inferred from endpoint assays like UV-Vis, HPLC, or mass loss. Affinity measurements are rarely reported. Instead, our understanding of these interactions is largely inferred from QM/MM and computational simulations using short oligomers that suggest nonspecific and relatively weak interactions.^15–18^ Supporting these findings, isothermal titration calorimetry (ITC) experiments carried out on the PDE *Tf*Cut2 show that 95% of degradation heat arises from cleavage, and only 5% from binding. These results suggested that interaction with PET is weak and catalysis may be diffusion-limited,^19^ although bias remains from reliance on an inactivated mutant that may possess a different surface affinity than the parent enzyme. Conversely, another study indirectly measuring depletion of PET hydrolases in solution by PET powder instead suggested a nanomolar binding affinity. However, these measurements required aqueous suspensions of highly hydrophobic particles that have historically led to shortcomings in standardization and reproducibility.^20^ This lack of consensus highlights the need for unbiased and standardizable methodologies to study PDE-plastic interaction.

To address this gap, we developed a simple, reproducible method to directly quantify PET hydrolase surface kinetics using surface plasmon resonance (SPR). SPR enables real-time, label-free tracking of enzymes binding to surfaces that no other method has achieved with comparable resolution.^21–23^ Adapting this approach to measure the binding of enzymes to PET surfaces, we measured nanomolar affinities and slow dissociation kinetics for four PET hydrolases. Strikingly, our data suggest that previously unreported structural rearrangements occur during PDE-plastic interaction in PET hydrolases. Moreover, binding disruption by degradation products such as terephthalic acid (TPA) and bis(2-Hydroxyethyl) terephthalate (BHET) supports a tight balance between priming surface binding and steric inhibitory effects. Overall, our results address uncertainty in the field regarding modes and affinities of PET hydrolase binding to authentic plastic surfaces. We find stable, long-lived, and dynamical enzyme-substrate interactions, suggesting that productive binding plays a more significant role in degradation than previously thought. By directly quantifying PET hydrolase kinetics on real plastic, we define unseen characteristics of PET recognition and offer valuable insight for future mechanistic studies and enzyme engineering.

## Results

### PET-coated SPR chips capture PET hydrolase-PET binding interactions

As SPR provides direct measurements of molecular binding near a plasmonic surface, we considered two architectures to study PDE-plastic interaction. Most SPR studies immobilize the protein to a microfluidic SPR chip before flowing a binding partner (**Figure 1a**). The capture and binding of biological constructs such as exosomes and even whole cells has been studied using such methods.^24^ However, plastic nanoparticles are highly prone to aggregation, deposition, and poor flow behaviour, and even the more tractable 3PET analog generated concentration-inverted and poorly reproducible SPR curves consistent with aggregation of the analog, preventing reliable kinetic analysis (**Supplementary Figure S1**). Therefore, we chose to instead immobilize a thin layer of plastic on the chip surface and flow a solution of enzyme to directly study enzyme binding to the modified chip surface. (**Figure 1b**)

**Figure 1.**
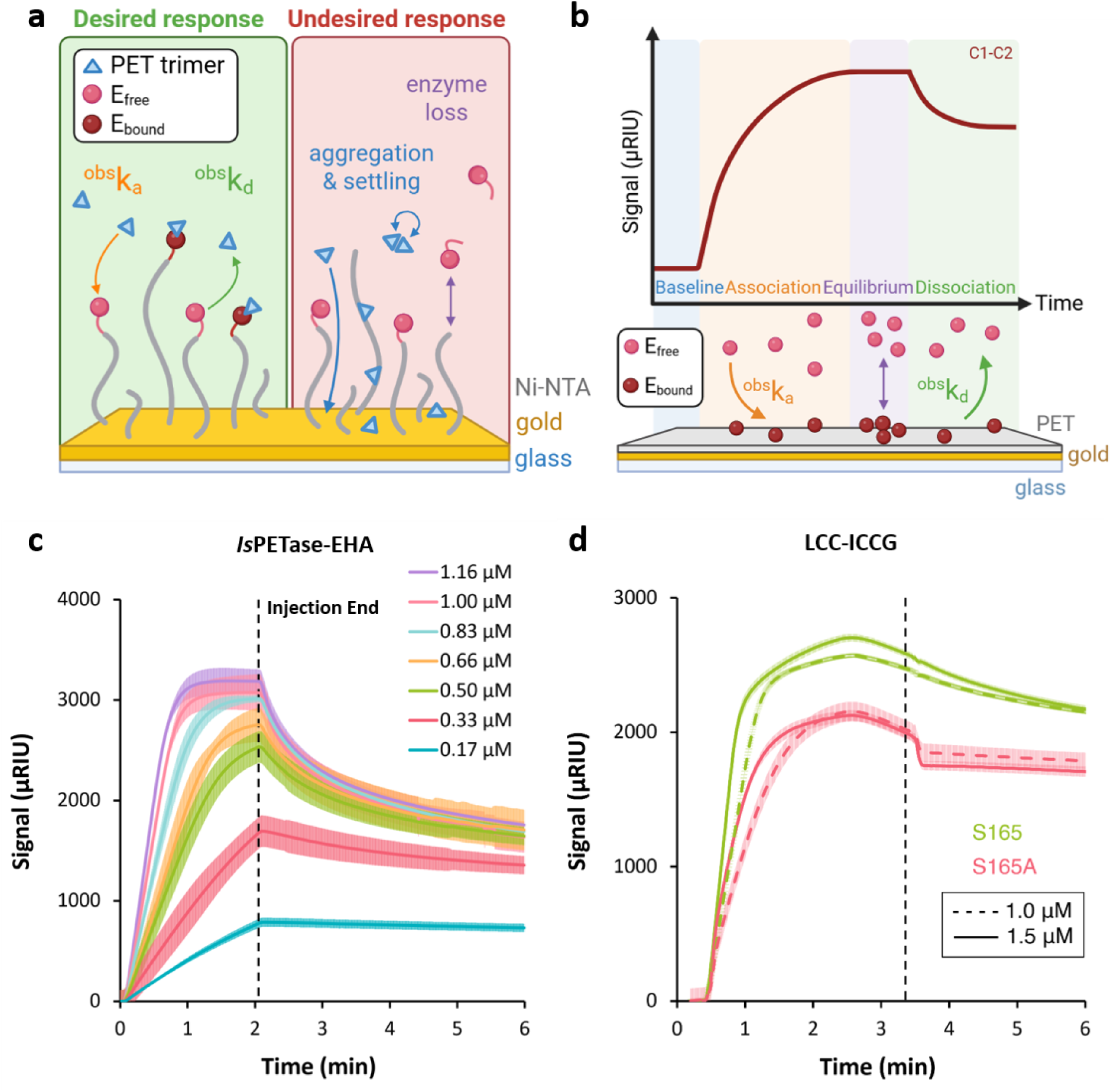
Direct measurement of PET hydrolase binding to PET-coated SPR surfaces. **(a)** Classical SPR paradigm. An enzyme is immobilised on a gold SPR chip and a analyte such as a suspension of 3PET flowed over the surface. In practice, plastic hydrophobicity complicates the flow regimes of suspensions due to noise-elevating processes such as aggregation and settling. (**b**) Inverted SPR paradigm used in this study. A thin amorphous PET film is deposited on a gold SPR chip and enzyme is flowed over the surface, allowing binding to be followed in real time. An idealised 1:1 response is shown, with the baseline, association, equilibrium and dissociation phases indicated. (**c**) Sensorgrams for 0.17–1.16 µM *Is*PETase-EHA injected over a PET-coated chip at 45°C in 20 mM HEPES, pH 8 at 25 µL/min. Dashed lines indicates the end of injection, which coincides with the maximum response. Shading indicates standard deviation across n = 3 injections. (**d**) Sensorgrams for LCC-ICCG (green) and its catalytically inactivated variant S165A (blue) at 1.0 µM (dashed) and 1.5 µM (solid), 45 °C in 20 mM HEPES, pH 8 at 25 µL/min. In both cases the response reaches a maximum at approximately 2.5 min, roughly one minute *before* the end of injection at 3.5 min, and declines while enzyme continues to flow over the surface. Retention of this behaviour in S165A establishes that it does not require catalytic turnover.

We used spin-coating to deposit a thin layer of PET on the surface of plain gold SPR chips. The resulting plastic-coated chip demonstrated a reduction in SPR trough intensity and a shift in the SPR detector scan of roughly 32 px, corresponding to the deposition of a PET layer of ∼10 nm thickness (**Supplementary Figure S2a**). Further validation of plastic deposition by atomic force microscopy showed a substantially different surface profile for PET-coated chips relative to bare gold chips, with a scratch test demonstrating a continuous soft layer with a thickness of ∼7 nm (**Supplementary Figure S2b,c**). Confocal Raman microscopy verified the film’s composition, with two characteristic PET peaks at ∼1615 and 1730 cm^-1^ confirming the presence of PET on the chips (**Supplementary Figure S2d,e**).^25^

Using these PET-modified chips, we ran SPR experiments to observe the binding of various PET hydrolases to a PET surface. Our initial enzyme test set included two of the most studied engineered PET hydrolases reported in the literature, leaf-branch compost cutinase (LCC)-ICCG and *Is*PETase-EHA, along with well-studied natural PDEs LCC and *Tf*Cut2.^1,26–28^ In all cases, ordered binding of PDEs to the PET chip surface was observed, with all four enzymes demonstrating reproducible, specific, concentration-and saturation-dependent binding (**Figure 1 cd**, **Table 1, Supplementary Figures S3, S4**), while a bare gold chip control showed weaker binding for all PET hydrolases (**Supplementary Figure S5**). Interestingly, while *Is*PETase-EHA displayed monophasic behaviour over the concentration range studied, LCC-ICCG and *Tf*Cut2 instead demonstrated multiphasic binding behaviour over long association periods. Initial linearity nonetheless allowed a good fit to the Langmuir 1:1 binding model and the determination of apparent binding kinetics for all enzymes except *Tf*Cut2, due to the latter’s strongly multiphasic character. All three other proteins demonstrated tight binding characteristics with nanomolar apparent affinities and long complex half-lives (>10 minutes) resulting from moderate apparent association rates (10^3^-10^4^ M^-1^s^-1^) and slow apparent dissociation rates (10^-4^ s^-1^). We also examined the binding properties of three non-PDEs: BSA, a non-specific hydrophobic binding protein, lysozyme, an enzyme known to bind tightly to PET,^29^ and *Pl*Cut, a non-plastic degrading cutinase with high sequence and structure similarity to the above PET hydrolases.^30^ These proteins either did not display specific or saturation-dependent binding, had poor reproducibility, and/or a U-value>10%, indicative of disordered binding modes.

**Table 1.**
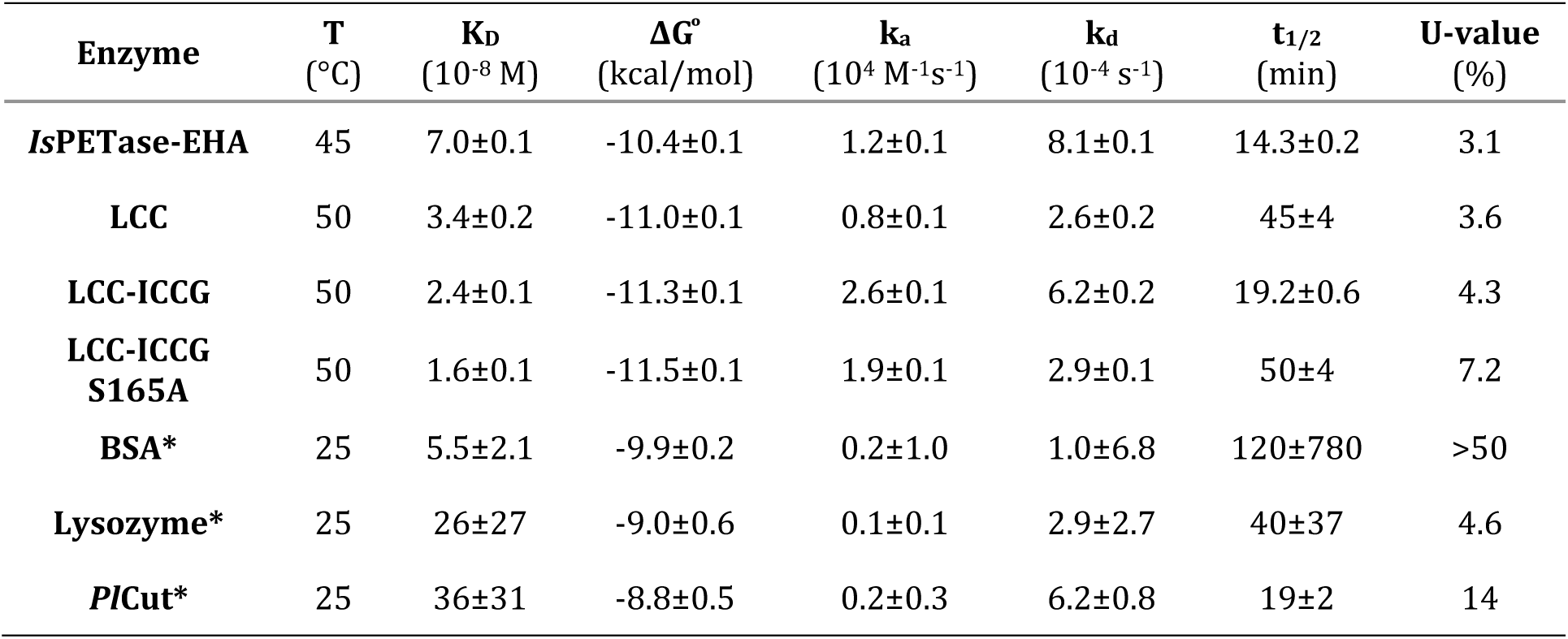
Apparent binding kinetics for PET hydrolases and non-degrading control proteins on PET-coated SPR chips. . Apparent equilibrium dissociation constant (K_D_), apparent Gibbs free energy of binding (ΔG ͦ), apparent association and dissociation rate constants (k_a_, k_d_), complex half-lives, and U-values (%) for protein-plastic interaction derived from SPR sensorgrams. Parameters were obtained by global fitting of a 1:1 Langmuir binding model to a concentration series of n = 3 replicate injections ± 1 SD. Assays were performed in 20 mM HEPES, pH 8. Enzymes marked (*) did not satisfy the fit acceptance criteria described in Materials and Methods, showing poor U-values and/or parameter uncertainties comparable to or exceeding fitted values. These are reported to document non-specific binding rather than as reliable kinetic constants.

### PETases binding is implicated in enzyme-dependent surface remodeling

The curvature observed in the sensorgrams of LCC-ICCG and *Tf*Cut2 is particularly striking given the reduction in SPR signal that is observed at the end of the association phase, reflecting a reduction in apparent surface-coupled mass. As macroscopic release of enzyme from the surface is unlikely during this phase, we considered possible mechanisms for this decrease. Given that PET hydrolases depolymerize PET, it is possible that plastic mass loss from enzyme catalysis is being observed. However, no decrease in the baseline signal was observed over the course of our experiments, thus indicating that no SPR-detectable damage to the plastic film is occurring (**Supplementary Figure S6**). In addition, a prior study of enzyme loading has shown that PET hydrolases likely form monolayers rather than layered aggregates on plastic surfaces that may induce complex binding behaviour.^20^ More strikingly, *Tf*Cut2 displays both the greatest deviation from classical kinetics and the lowest peak signal, further suggesting that this unusual curvature is not the result of multi-layered binding. Increasing temperature gradually eliminated the SPR signal reduction (**Figure 2 ab**), further confirming that this effect is not plastic loss, which would be enhanced at high temperature. Rather, at elevated temperature, the rearrangement becomes fast relative to association and is no longer resolved as a distinct phase. As a result, the decline in response most plausibly reflects a slow reorientation or conformational change in the enzyme and/or surface that moves the complex center of mass farther from the surface, decreasing its effective signal contribution. The instrument used to generate these results is prism-based, with *l*d ≈ 200 nm. Producing a decline of ≥ 10 % of the association signal by alteration of the enzyme layer alone would require its mass centroid to move by roughly 10 nm away from the surface. PET hydrolases are approximately 4 nm in diameter, so no enzyme conformational change can satisfy this requirement alone. This signal change therefore must implicate the polymer, and the redistribution of a small fraction of the film to a 10 nm distance is consistent with both our observations and polymer chain lengths for virgin commercial PET, which routinely exceed 100 nm. It is therefore likely that the signal drop reports an enzyme-dependent surface remodelling effect.

**Figure 2.**
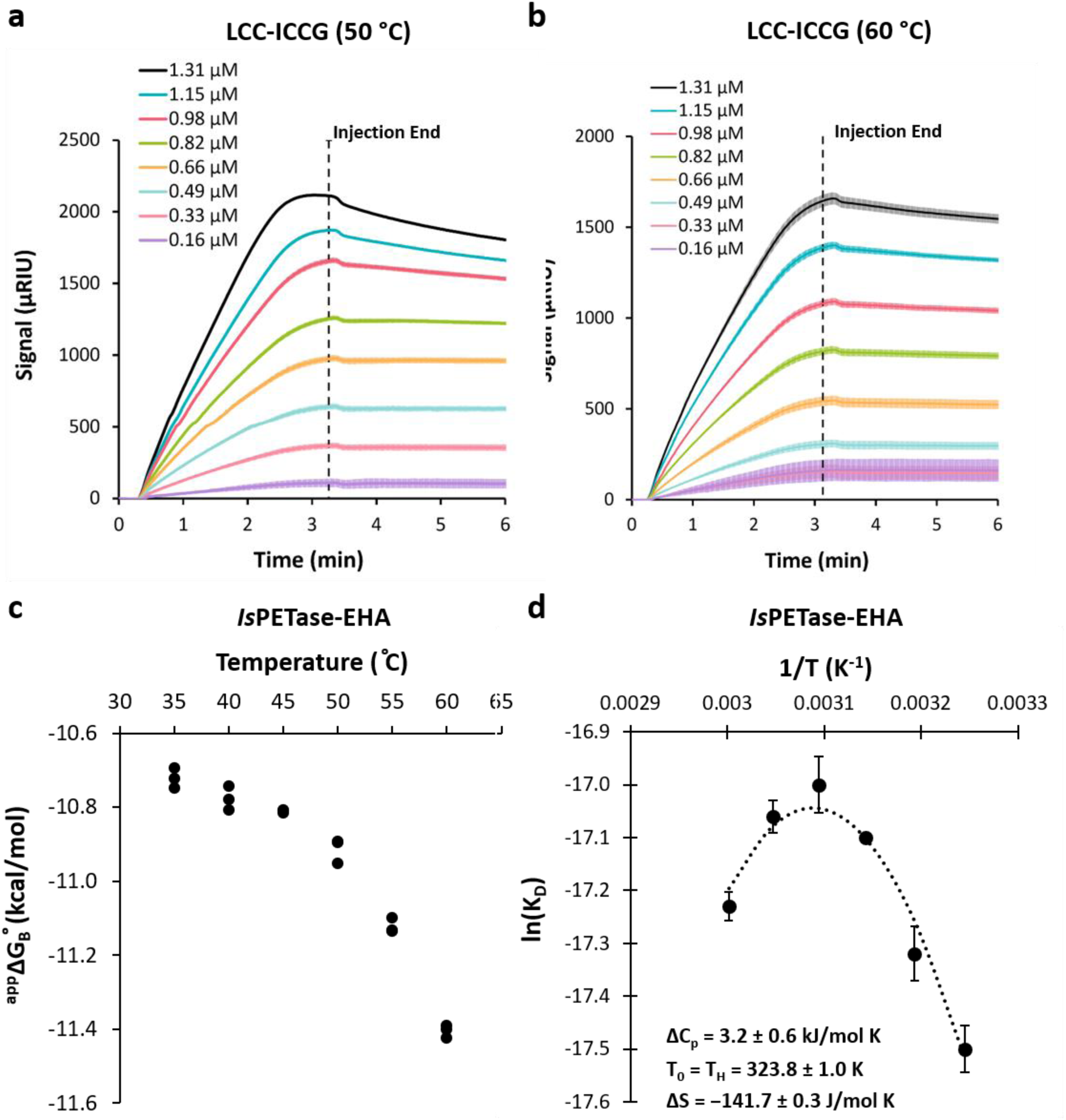
Temperature dependence of PET hydrolase surface engagement. (**a, b**) SPR sensorgrams for 0.16–1.31 µM LCC-ICCG injected over PET-coated chips at (a) 50 °C and (b) 60 °C in 20 mM HEPES, pH 8 at 25 µL/min. Dashed lines indicate the end of injection. Shading indicates standard deviation across n = 3 injections. (**c, d**) Temperature dependence of *Is*PETase-EHA binding to PET, measured from 35 to 60 °C (n = 3 ± 1 SD). (**c**) Apparent Gibbs free energy of binding (^app^ΔG°_B_) becomes progressively more favourable with increasing temperature. (**d**) The corresponding van’t Hoff plot is markedly non-linear, with apparent *K*_D_ weakest near 50 °C and tighter at both temperature extremes. Fitting the generalised van’t Hoff equation with a temperature-dependent enthalpy correction (dotted line, *T*₀ = *T*_H_ = 323.8 ± 1.0 K) gives ΔC*_p_* = 3.2 ± 0.6 kJ mol⁻¹ K⁻¹ and ΔS° = −141.7 ± 0.3 J mol⁻¹ K⁻¹. A positive heat capacity change of this magnitude is inconsistent with rigid-body adsorption and indicates that binding is accompanied by hydrophobic surface exposure.

Unlike LCC-ICCG, *Is*PETase-EHA presented classical monophasic response curves, implying that this enzyme does not undergo a resolvable conformational rearrangement upon binding to the PET surface. However, this does not discount the possibility of a rearrangement occurring at a faster timescale than association, leading to an apparent monophasic process in single SPR curves. Therefore, to better understand this response, we evaluated the temperature dependence of *Is*PETase-EHA binding affinity (**Figure 2 cd**). The resulting ^app^ΔG_B_^°^ /T and van’t Hoff plots show clear non-linearity, with strong downward curvature indicating a positive ΔC_P_^app^ for this binding process. Fitting the generalized van’t Hoff equation with temperature-dependent enthalpy correction, we find an approximate ΔC_P_^app^ of +3.2 kJ/mol K. As a positive ΔC_P_ is related to hydrophobic surface exposure during the transition, this behaviour would not be expected for simple enzyme-plastic binding, but is consistent with the chain loosening effect hypothesized to occur during LCC-ICCG binding. Furthermore, apparent temperature effects on *K*_D_ are primarily related to changes in *k*_d_ rather than *k*_a_, indicating that temperature-driven changes in surface accessibility, which primarily affect *k*a, are not the causal effect of this behaviour (**Supplementary Figure S7**). Together, these results imply that a PET hydrolase-driven surface remodelling occurs following *Is*PETase binding as well. The accelerated rate for this process relative to LCC-ICCG is consistent with the greater structural flexibility and more open active site of *Is*PETase-EHA (**Figure 3, Supplementary Figure S8**).^31–33^

**Figure 3.**
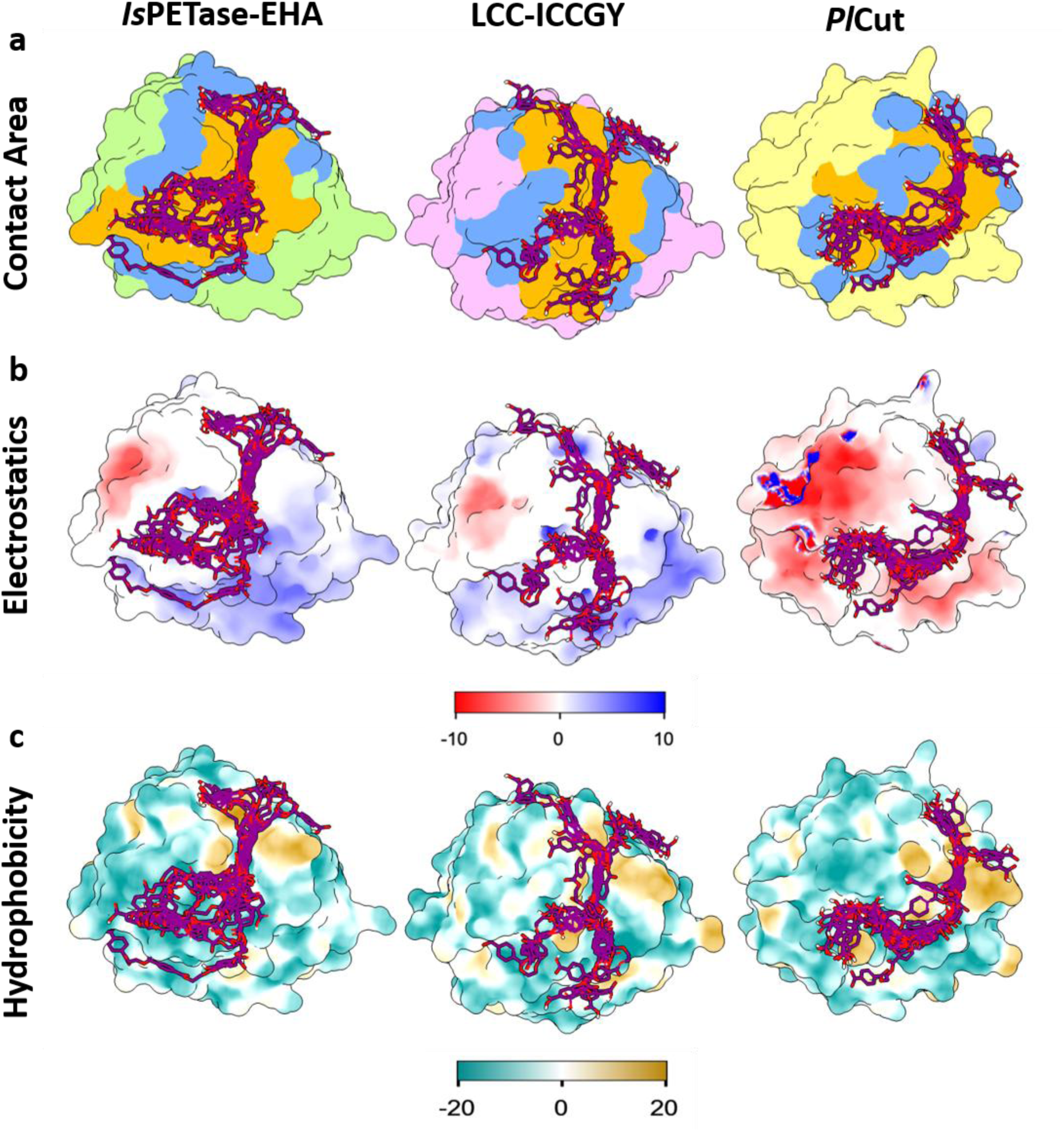
Structural comparison of the substrate-binding surfaces of PET-active and PET-inactive α/β-hydrolases. (**a**) Surface representations of *Is*PETase-EHA (PDB 6IJ6), LCC-ICCGY (PDB 8QRJ) and an AlphaFold3 model of *Pl*Cut, showing the canonical substrate-binding cleft (orange) and the surface contacted by a docked 4PET chain (blue). Docked poses (15-20 modes retained per enzyme) are shown in magenta. Contacted surface, cleft surface, and the fraction of contact falling outside the canonical cleft are given in Table SX. (**b**) Electrostatic potential mapped onto the solvent-accessible surface, calculated with APBS at pH 7.0 and 150 mM ionic strength, coloured from −3 to +3 k*T*/e (red to blue). (**c**) Surface molecular lipophilicity potential coloured from hydrophilic (teal) to hydrophobic (gold). Docked 4PET poses are shown in (b) and (c) for orientation. All structures are shown in the same orientation, with the substrate-binding cleft facing the viewer. Despite comparable cleft geometry, electrostatic and hydrophobic surface properties do not distinguish the two PET-active enzymes from *Pl*Cut, which does not productively engage PET under the conditions used here.

### Catalytic knockouts and common solubilizers bias binding kinetics

To establish that this observed behaviour is due to catalysis-independent surface engagement rather than damage to the plastic layer from enzyme activity, we recorded PET binding curves for the LCC-ICCG catalytic knockout S165A (**Figure 1d**). The downwards curvature was retained, confirming that the multiphasic binding curves observed are indeed a function of binding rather than catalysis. However, this mutant displayed significantly slower kinetics, with a *k*_d_ more than two-fold lower than the parent LCC-ICCG. In retrospect, this is not unusual, given that the increased hydropathy of alanine relative to serine (+2.6) would enable a tighter-binding engagement of highly hydrophobic PET strands. However, this result highlights that catalytic knockouts cannot be assumed to possess identical binding properties to their parents, leading to biases when used as non-catalytic binding energy controls.

This realization prompted us to further examine biases in common methods of studying PET hydrolase-plastic interactions. Importantly, many studies that attempt to characterize these interactions use small molecule analogs like 3PET or suspensions of plastic nanoparticles. However, due to the aggregation propensity of these hydrophobic analogs, maintaining these suspensions commonly requires the use of stabilizers such as Triton X-100 (TX100) or polyvinyl alcohol (PVA).^34–36^ As these stabilizers interact directly with both the plastics and enzymes, it is reasonable to expect that they would alter protein-plastic interactions. To quantify these biases, we recorded PET binding curves for both LCC-ICCG and *Is*PETase-EHA in the presence of TX100 (**Supplementary Figure S9**). Interestingly, while increasing concentrations of TX100 mildly reduced enzyme-surface affinity, a more dramatic effect on curve shape was observed. For LCC-ICCG, the curvature that had been kinetically collapsed by high temperature was gradually restored, indicating that these stabilizers decelerate this rearrangement. This effect could be observed at TX100 concentrations as low as 0.005 %v/v, with full downwards curvature returning at TX100 concentrations of 0.05 %_v/v_ and above. Similarly, a slight downward curvature developed in *Is*PETase-EHA association curves, most visibly at 0.1 %_v/v_, which may suggest that rearrangement is being slowed towards resolvability. In both cases, the TX100 concentrations at which these kinetic effects are observed are representative of the 0.01 – 0.1 %_v/v_ concentration range generally used in characterization experiments.

### PET hydrolysis products modulate surface engagement

As PET hydrolases are widely reported to be product inhibited, we next evaluated the effects of the PET degradation products TPA and BHET on enzyme-plastic binding affinity (**Figure 4a-c, Supplementary Figure S10**). As stabilizers are needed to flow these products, we validated these results with both TX100 and PVA alongside a no-inhibitor control with added stabilizer. Similar trends were observed for both *Is*PETase-EHA and LCC-ICCG. However, strong stabilizer-dependent effects were seen. Although TPA and BHET have been thought to act competitively, we observed that at low concentrations and in the presence of 0.01% PVA as a stabilizer, they instead enhance surface occupancy for both LCC-ICCG and *Is*PETase-EHA, as evidenced by an increased peak signal without altering binding kinetics. As product concentration increases further, however, a progressive reduction in surface accessibility can be observed, ultimately overtaking the initial enhancement and leading to overall inhibition of surface engagement by PET hydrolases. These effects were however not observed with 0.01% TX100, with all product curves remaining superimposable for both enzymes.

**Figure 4.**
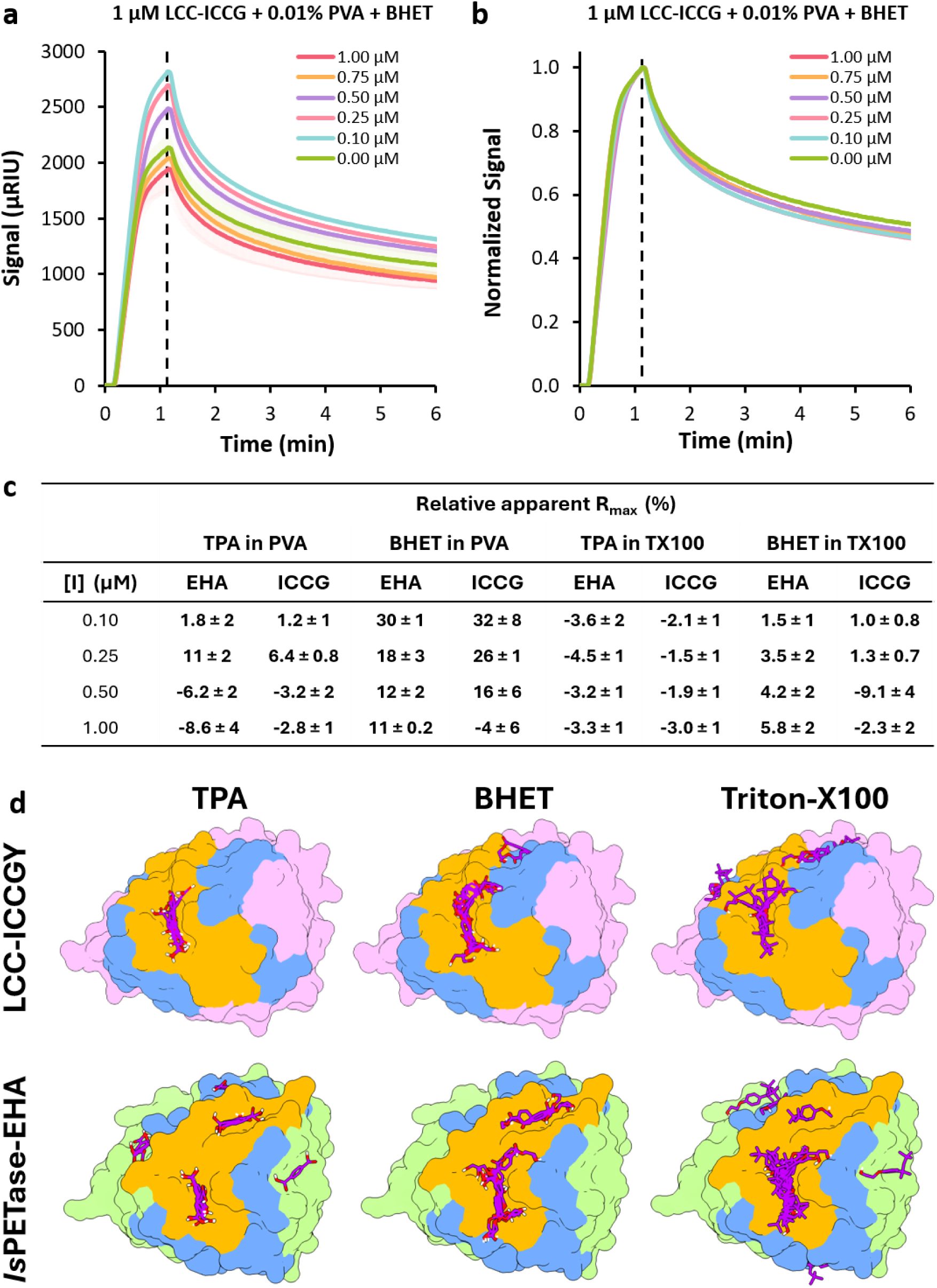
PET degradation products modulate the extent of PET hydrolase surface engagement. (**a**) Sensorgrams for 1 µM LCC-ICCG injected over a PET-coated chip in 20 mM HEPES pH 8 with 0.01% PVA, in the presence of 0–1.00 µM BHET, at 60 °C and 25 µL/min. The dashed line indicates the end of injection. Shading indicates standard deviation across n = 3 injections ±1 SD. (**b**) The same sensorgrams normalised to their individual maxima. Association and dissociation kinetics are superimposable across the series, indicating that BHET alters the extent of enzyme accumulation on the surface rather than the rate constants describing it. (**c**) Apparent Rmax values from fitted sensorgram curves for 0.10 – 1.0 μM BHET and TPA injected with 1 μM *Is*PETase-EHA or LCC-ICCG over a PET-coated chip in 20 mM HEPES pH 8 with 0.01% PVA or TX100, at 45 °C (*Is*PETase-EHA) or 50°C (LCC-ICCG) at 25 μL/min across n = 3 injections ± 1 SD. (**d**) Representative docked poses of TPA, BHET and Triton X-100 in *Is*PETase-EHA (PDB 6IJ6, top) and LCC-ICCGY (PDB 8QRJ, bottom). The catalytic triad is shown in yellow. Poses were used to assess positional accommodation within and around the binding cleft. All three ligands are accommodated at overlapping positions in the cleft of both enzymes.

Two potential hypotheses explain this behaviour. TPA and BHET are both plausible plasticizers of polyesters ^37^ and may expose additional PET hydrolase binding sites by partitioning into the surface and loosening strands. However, no baseline shifts are observed between experiments, indicating that these effects are transient. Additionally, given the multivalency of PET hydrolase binding sites,^38–40^ it is possible that partial binding cleft occupancy may prime the enzyme for higher affinity surface interaction, while higher-valency occupancy at high product concentration instead promotes inhibition. TX100, meanwhile, is also capable of acting as a PET plasticizer, and docking simulations to LCC-ICCG and *Is*PETase-EHA reveal similar interactions for TPA, BHET, and TX100 with PET hydrolases (**Figure 4d**). As a result, as TX100 concentrations far exceed those of TPA and BHET used, it is possible that the use of TX100 masks the effects of these products, while PVA, a steric stabilizer that is not predicted to plasticize PET or interact meaningfully with PET hydrolases, does not.

## Discussion

Efforts to engineer PET-degrading enzymes have overwhelmingly targeted thermostability and active-site chemistry. Our results indicate that the capacity to engage and mobilise polymer chains at the interface is distinct from both and largely invisible to many assays through which the field measures success. The PET hydrolases examined here bind PET with nanomolar apparent affinities and dissociate slowly, with apparent surface residence half-lives measured in tens of minutes. These values are at odds with turnover numbers reported by many analog-based assays for the same enzymes. A recent study using a 3PET substrate reported an LCC k_cat_ of 4.04 s^-1^.^41^ However, if applied to PET, our measured LCC k_d_ of 2.3∗10^-4^ s^-1^ would imply approximately 2∗10^4^ cleavages per encounter to achieve a similar k_cat_. Processivity on that scale is not supported by the structure of these enzymes, which possess open, solvent-exposed clefts. Conversely, inverse Michaelis-Menten kinetics performed on authentic PET for LCC-ICCG report a k_cat_ of 7.8∗10^-4^ s^-1^.^42^ This value is strikingly similar to our measured k_d_ of 4.0∗10^-4^ s^-1^, which, alongside the 3PET data, supports that turnover in these enzymes is limited by physical interaction with the plastic surface and that ester bond cleavage occurs considerably faster than disengagement. By this logic, the modest catalytic efficiency of PET hydrolases relative to their effectiveness on PET is not paradoxical, but a direct consequence of interfacial kinetics, in which the limiting event is the acquisition, maintenance, and release of a productive enzyme-chain complex rather the hydrolysis step itself.

What occurs during this residence time is directly visible in our data. The signal reduction observed during enzyme flow for *Tf*Cut2 and LCC-ICCG, non-monotonic temperature dependence of *Is*PETase binding, and deviation from Langmuir one-to-one binding curve shapes of LCC all point towards complex enzyme-surface interaction. Strikingly, the signal reduction is retained in the catalytically inactivated S165A variant, while being absent in BSA, lysozyme, and the structurally homologous but non-plastic-degrading cutinase *Pl*Cut. We interpret this as an enzyme-dependent mobilisation of surface chains, comparable in effect to the non-hydrolytic disruption of cellulose chains long recognised in cellulose-degrading systems.^43^

Although SPR-observable effects are likely tied to polymer chain mobility, LCC-ICCG and *Tf*Cut2 resolve the process to a directly observable extent over 3 minutes, LCC does not do so within our injection window despite comparable early association behaviour, and *Is*PETase-EHA appears monophasic, consistent with a rearrangement faster than association. Given that the architecture of our PET-coated chips is common to all experiments, these different kinetics regimes cannot arise from the intrinsic properties of the film. These results also have implications for how the temperature dependence of these enzymes is usually explained. Enhanced activity at elevated temperature is commonly attributed to increased chain mobility as PET nears its glass transition temperature (*Tg* = 78 °C); however, the estimated hydrated *Tg* of PET ( ≈ 60 °C) and the surface-layer *Tg* ( ≈ 40-45 °C) are both substantially lower than the optimum operating temperatures of thermostable PET hydrolases such as LCC-ICCG.^44^ Furthermore, it was recently shown that depressing substrate *Tg* by pre-soaking did not enhance LCC-ICCG activity.^45^ Here the same film is remodelled at different rates according to which enzyme is present. Reconciling the above, we suggest that the dominant temperature-dependent variable for LCC-class enzymes, and perhaps more broadly for PET hydrolases, appears to be at least partially related to conformational sampling by the enzyme.

By this view, LCC-derived and *Is*PETase-derived enzymes appear to solve the same interfacial problem by different routes. *Is*PETase-EHA embeds flexibility and aromatic stacking directly into its cleft, leading to unresolvable remodelling on our measurement timescale. The functional importance of this architecture is well established, since converting the *Is*PETase cleft towards a *Tf*Cut2-like geometry abolishes more than 90% of PET hydrolysis,^46^ and substituting disulfides with cutinase-derived residues is similarly deleterious to catalysis.^31,47^ LCC-class cutinases instead present a more rigid ground-state cleft and correspondingly slower remodelling, which accelerates with temperature until it can no longer be resolved kinetically. This behaviour is consistent with temperature-dependent access to conformations that *Is*PETase-EHA readily accesses at ambient temperature. Most interestingly, this ordering tracks reported activity trends, albeit over a small number of enzymes.

Extending these surface remodelling effects to the concept of a Sabatier optimum, it is important to contrast the different interactions that occur during surface engagement. An enzyme may bind the polymer tightly without extracting a chain, and conversely, an enzyme that extracts chains efficiently may not release them readily afterwards. The relevant measurement is therefore not solely the affinity of the enzyme for the polymer surface, but rather its partition between surface association and chain mobilisation. Two independent lines of evidence support this. First, surface residence studies report that PET hydrolases with high activity show higher protein motility, indicating that residence time and activity are not monotonically related.^10,48,49^ Second, engineering LCC to reduce PET binding affinity improved product yield with F243I, a substitution also present in LCC-ICCG, decreasing hydrophobic surface interaction and allowing faster product release. However, F243W, a cleft-narrowing substitution at the same position, also improved conversion, highlighting that these effects are not reducible to a simple two-state affinity coordinate.^50^

Degradation products modulate this balance rather than simply opposing it. Sub-inhibitory concentrations of TPA and BHET increase surface engagement, with inhibition emerging only above approximately 0.75 μM. Transient partitioning into the film, increasing local chain mobility, and partial occupancy of the cleft favouring a more engagement-competent conformation both remain consistent with these data. While we cannot presently distinguish between these processes, either extends rather than contradicts the established picture of PET degradation products as competitive inhibitors of PET hydrolases.^51,52^ Because local product concentration at the interface must necessarily pass through the activating regime before reaching inhibitory levels, it may contribute to the non-linear progress curves widely reported for these enzymes.

Furthermore, if chain mobilisation is a distinct, rate-limiting step dependent on enzymatic conformational accessibility, several persistent problems in engineering these enzymes becomes more understandable. Ancestral reconstruction has shown that PET hydrolase activity arises along multiple independent trajectories involving distal mutations, across a highly rugged landscape where phylogenetically equivalent nodes differ substantially in activity.^8,9^ This observation is difficult to rationalize if function is tied to active-site chemistry and thermostability but expected if the limiting step is a dynamical property sensitive to mutations throughout the fold. The same logic accounts for an oft-seen artifact in PET hydrolase engineering, with variants that are successfully stabilized and improved as esterases often displaying diminished activity versus PET. Stabilization generally proceeds by fold rigidification, and if chain extraction requires a specific dynamical trajectory that is not resolved in known PET hydrolase crystal structures, rigidifying the fold removes this capability while leaving chemistry intact. This loss is invisible to analog-based assays and is therefore usually detected late. It further suggests that thermal robustness and conformational access are separable engineering axes, only one of which has been systematically exploited.

Two features of our approach limit our conclusions. Our PET-coated chips are spin-coated with highly pure amorphous PET of approximately 10 nm thickness and differ from post-consumer material in crystallinity, thickness, and additive presence. The values we report should therefore be viewed as a characteristic of this substrate and may be only partially transferrable. We also measure interfacial engagement rather than turnover, with the correspondence between them inferred through exclusion of confounding hypotheses rather than conclusively demonstrated. Direct observation of the mobilised layer remains to be performed. Nonetheless, our ability to exclude biasing factors such as stabilizers leaves these measurements as some of the most direct analyses of PET hydrolase interactions with authentic plastic surfaces reported to date. This may explain why these effects have not previously been reported, since they are suppressed by the dilute TX100 and PVA required to maintain nanoparticle and small molecule suspensions on which much of the field’s interaction data rest.

Overall, by measuring PET hydrolase binding directly on authentic PET, we find that these enzymes engage their substrate tightly, persistently, and in a manner that reshapes the polymer surface. Depolymerisation of PET therefore involves at least one step beyond ester hydrolysis, the mobilisation of a chain from a condensed phase, that no soluble analog can report and that current assays are not designed to detect. Recognising this step as distinct and potentially limiting offers a route to understanding why these enzymes have proven so resistant to rational improvement, and a concrete target for the improvement itself.

## Materials and Methods

### Materials

#### Plastic & monomers

300 μm PET powder of >50% crystallinity (ES30-PD-000132) and 0.1 mm thick PET film (ES30-FM-000200) were purchased from Goodfellow. Bis(2-hydroxyethyl) terephthalate (BHET) was purchased from TCI America (B3429) and terephthalic acid (TPA) was purchased from Thermo Scientific (180725000). 3PET was gifted from the Howe Lab (Queen’s University) *Plasmids:* Genes encoding *Is*PETase-EHA, LCC, LCC-ICCG, *Tf*Cut, and *Pl*Cut were synthesized in pET29b vectors by Twist Bioscience. A catalytic knockout of LCC-ICCG, LCC F243I/D238C/S283C/Y127G/S165A, was synthesized by KLD treatment according to NEB (#M0554) and sequenced by nanopore sequencing. *Lyophilized enzymes:* Lysozyme (100831 MP biomedicals) and BSA (A7030-10G MilliporeSigma) were dissolved in 20 mM HEPES pH 8 prior to use.

### Preparation and Characterization of PET-coated SPR Chips

#### Amorphous films

Gold 1×1cm SPR chips (Reichert) were spin coated with 1-5% PET powder dissolved in +99% TFA at 2500 rpm for 30 seconds. Spins were repeated with +99% TFA aliquots until the surface appeared homogenous and produced equivalent C1/C2 detector scan peaks. *Characterization*: Chips were evaluated by SPR pre-and post-lamination to estimate ΔμRIU. A neat gold and PET chip were characterized by Raman spectroscopy and atomic force microscopy (AFM) to determine the film’s composition and thickness. Raman spectroscopy measurements were taken using a Renishaw in Via Qontor confocal Raman microscope that uses a Leica Microsystems bright-field microscope with a DM2700 light source. A 500 mW 532 nm wavelength laser with a 2400 L mm^−1^ grating was used to obtain measurements in the spectral range of 1100– 2000 cm^−1^, focused on the sample by an X50L objective. The spectra were acquired with 50% laser power and 1 second exposure time. AFM images are taken using a Bruker Dimension FastScan AFM with ScanAsyst-Air tips, running 512 scan lines in PeakForce Tapping mode. Images were then processed with Nanoscope Analysis.

### Recombinant Protein Expression & Purification

#### Recombinant protein expression

PDEs were expressed in LOBSTR BL21 *E. coli* before IPTG induction at 16 ͦC overnight. Cultures were centrifuged at 6000g, 4 ͦC for 30 minutes (CR22N Himac) before pellets were resuspended in equilibration buffer on ice (100 mM sodium phosphate buffer pH 7.4, 150 mM NaCl, and 10 mM imidazole). *Protein extraction & purification*: 10 mg lysozyme (Thermo Fisher Scientific) and 250U benzonase nuclease (Sigma Aldrich) were added to the cell suspension before 3 minutes of sonication for 10 seconds on, 10 seconds off, at 50% power on ice. Samples were centrifuged at 16000g, 4 ͦC for 30 minutes and the supernatant was passed through a 0.45 μm PVDF syringe filter (Sartorius). The clarified sample was applied to a column of HisPur Ni-NTA resin (Thermo Fisher), washed with equilibration buffer supplemented with 60 mM imidazole, and eluted in buffer containing 300 mM imidazole. The eluent was spin concentrated in a 2 kDa Vivaspin 15R column (Sartorius) before FPLC purification in assay buffer (Superdex 75 10/300 GL; AKTApure, Cytiva) at 4 ͦC. Fractions of the pellet, clarified sample, column flowthrough, column wash 1/2, final elution, and FPLC samples were collected for visualization by SDS-PAGE (12%, 225V, 45 min; Thermo Scientific PageRuler plus).

### SPR with PET-coated chips

#### Experimental

SPR was performed on the 2SPR Reichert system equipped with a 100 μL injection syringe. Channel 1 (C1) is designated for kinetics and channel 2 (C2) is designated as the control. Enzyme aliquots were injected over 1-3 minutes with a 10-20 minute dissociation period followed by surface regeneration with 0.1 M NaOH. Enzyme concentrations were optimized in triplicate experiments at 25 μL/min. 20 mM HEPES pH 8 was used as the running buffer and blank injections were performed before each enzyme injection. Lysozyme/BSA/*Pl*Cut were evaluated at 25 ͦC, *Is*PETase-EHA was evaluated at 45 ͦC, and *Tf*Cut and LCC variants were evaluated at 50 OR 60 ͦC. *Inhibition assays: Is*PETase-EHA and LCC-ICCG were assessed in the presence of 0.01-1.0 μM TPA/BHET and, for LCC-ICCG, 0.01-1.0 μM 3PET. Stock TPA/BHET/3PET solutions were prepared in 0.1-1% Triton-X100 (9410-1L MilliporeSigma) or PVA (9,000-10,000 g/mol; 360627-25G Sigma Aldrich) before being diluted in running buffer to 0.01-0.05%. Similarly, 20 mM HEPES pH 8 with 0.01-0.05% Triton-X100 or PVA was used as the running buffer. 1.0 μM *Is*PETase-EHA and LCC-ICCG were assessed at 45 ͦC and 60 ͦC, respectively (n=3). *Temperature dependence of apparent binding affinity:* 1.0 *μ*M aliquots of *Is*PETase-EHA in 20 mM HEPES pH 8 were injected at 35, 40, 45, 50, 55, and 60 *֯* C (25 uL/min, n=4). 1.0 – 1.5 *μ*M aliquots of LCC-ICCG and LCC-ICCG S165A were injected at 45, 50, and 60 *֯* C (25 uL/min, n=3). *Analysis*: SPR signal (μRIU) vs time curves were collected in TraceDrawer and C2 subtraction was performed. Buffer injections executed before enzyme injections were subtracted from the latter to further correct for buffer mismatch. Kinetic parameters were determined using a 1:1 Langmuir binding model with local (reported for figures) and global (reported for kinetic parameters) fitting. A fit was deemed acceptable if (1) full baseline regeneration was achieved, (2) the corresponding U-value was <10%, and (3) the association rate scaled linearly with the concentration of enzyme until saturation is reached. Kinetic parameters for comparison between two groups were log-transformed and analysed by Welch’s t-test (two-tailed, unequal variance, α=0.05). For assays evaluating changes in kinetic parameters in the presence and absence of inhibitors, data was log-transformed before ANOVA single factor and Tukey HSD analysis (α=0.05).

### SPR with a PET Trimer and His-Tag Immobilized PETases

#### Poly His-tag immobilization

Nickel-nitrilotriacetic acid SPR chips (Reichert) were prepared by injection of 200 mM NiCl_2_ over both channels. 100 μL of 0.5 *μ*M poly His-tagged enzyme was injected under the same conditions over C1 only. 3PET was dissolved in running buffer for injection of 100 μL over both channels. The running buffer was similarly diluted for buffer control injection; both the analyte and buffer samples were injected over 50 seconds to 3 minutes with a 10–15-minute dissociation. The surface was regenerated by injection of 200 mM imidazole followed by 60 mM EDTA over both channels. *IsPETase-EHA:* 0.05-0.5 *μ*M 3PET injected for 3 minutes at 25 uL/min, 40 *֯* C in 20 mM HEPES pH 8 0.01% Triton-X100. *LCC:* 0.01-0.2 *μ*M 3PET injected for 50 seconds at 45 uL/min, 50 *֯* C in 20 mM HEPES pH 8 0.01% Triton-X100. *LCC-ICCG:* 0.1-2.0 *μ*M 3PET injected for 3 minutes at 25 *μ*L/min, 60 *֯* C in 20 mM HEPES pH 8 0.05% Triton-X100. *Analysis*: As described previously.

### Computational structure assessment

#### Structure files

The x-ray crystal structures of PDB entries 6IJ6 (1.95 Å) and 8QRJ (1.42 Å; LCC-ICCGY) were used to represent *Is*PETase-EHA and LCC-ICCG, respectively. Crystal structures were stripped of crystallographic waters, ions, and bound ligands, hydrogens were added, and structures were protonated at pH 7 prior to docking (PyMOL v3.1.0, ChimeraX v1.10). Structure models for TPA, BHET, and Triton-X100 were downloaded from the ZINC database and 4PET was built in Avogadro v1.2.0 and parameterized using CGenFF v2.5.2, with rotatable bonds retained for flexible docking. *Docking:* A 50×40×96 Å search box surrounding the catalytic serine was used for TPA/BHET/Triton-X100 docking with AutoDock vina v1.1.2 (exhaustiveness=40, modes=15). For docking with 4PET, search box dimensions were adjusted to encompass the substrate-binding cleft of each enzyme (50×40×60 Å for 8QRJ and 36×32×62 Å for 6IJ6; exhaustiveness=40, modes=15). *Analysis:* Enzyme structure figures were prepared in ChimeraX. Electrostatic surfaces were calculated using the APBS plugin (v3.4.1; ionic strength of 150 mM, pH 7.0) in PyMOL and surface hydrophobics were calculated using Kyte-Doolittle hydrophobics (ChimeraX) before comparison with the webPIPSA server. SASA was calculated in PyMOL using the Shrake-Rupley algorithm and in ChimeraX using the rolling-probe algorithm. Enzyme pockets were assessed using the CASTpFOLD server. *Caver tunnel and MDS analysis*: Potential access tunnels in 6IJ6 and 8QRJ were analyzed using CAVERWeb. For 6IJ6, tunnel calculations were initiated from residues 97, 159, 182, 183, 201–203, 230, 232, 237–241, and 249–250, whereas for 8QRJ, starting-point residues were 93–97, 101–102, 164–165, 212, 242–243, and 246. For both structures, CAVER calculations used a probe radius of 0.9 Å, shell depth of 4.0 Å, shell radius of 3.0 Å, clustering threshold of 3.5 Å, maximum distance of 3.0 Å, and desired tunnel radius of 5.0 Å. Tunnel residues were computed, with a tunnel visualization sampling step of 0.5 Å, and a random seed of 1 was used for reproducibility. Dynamic tunnel visualization and trajectory generation were enabled. The resulting tunnels were subsequently evaluated using the associated YASARA molecular dynamics simulations. Simulations were performed at a temperature of 298.15 K and pH 7.4, with a system density of 0.997 g cm⁻³, for a total duration of 5 ns. Coordinates were saved at 10 ps intervals, resulting in 501 snapshots for each trajectory.

## Acknowledgements

We thank the Howe Group at Queen’s University for providing the 3PET used. We also thank the Scaiano and Lessard research groups at the University of Ottawa for use of their spin coating instrument. This research was enabled in part by the Digital Research Alliance of Canada.

## Contributions

KJP and AMD initiated and designed the project. LED and BHL designed and executed chip characterization experiments and analyzed the data. AB subcloned and transformed plasmids with catalytic knockouts. KJP and AMD designed and KJP carried out all other experiments and analyzed the data. KJP and AMD wrote the paper with feedback from all authors. AMD supervised the project.

