## Supplementary Data for "Direct measurement of PET hydrolase interfacial kinetics reveals catalysis-independent surface remodelling"

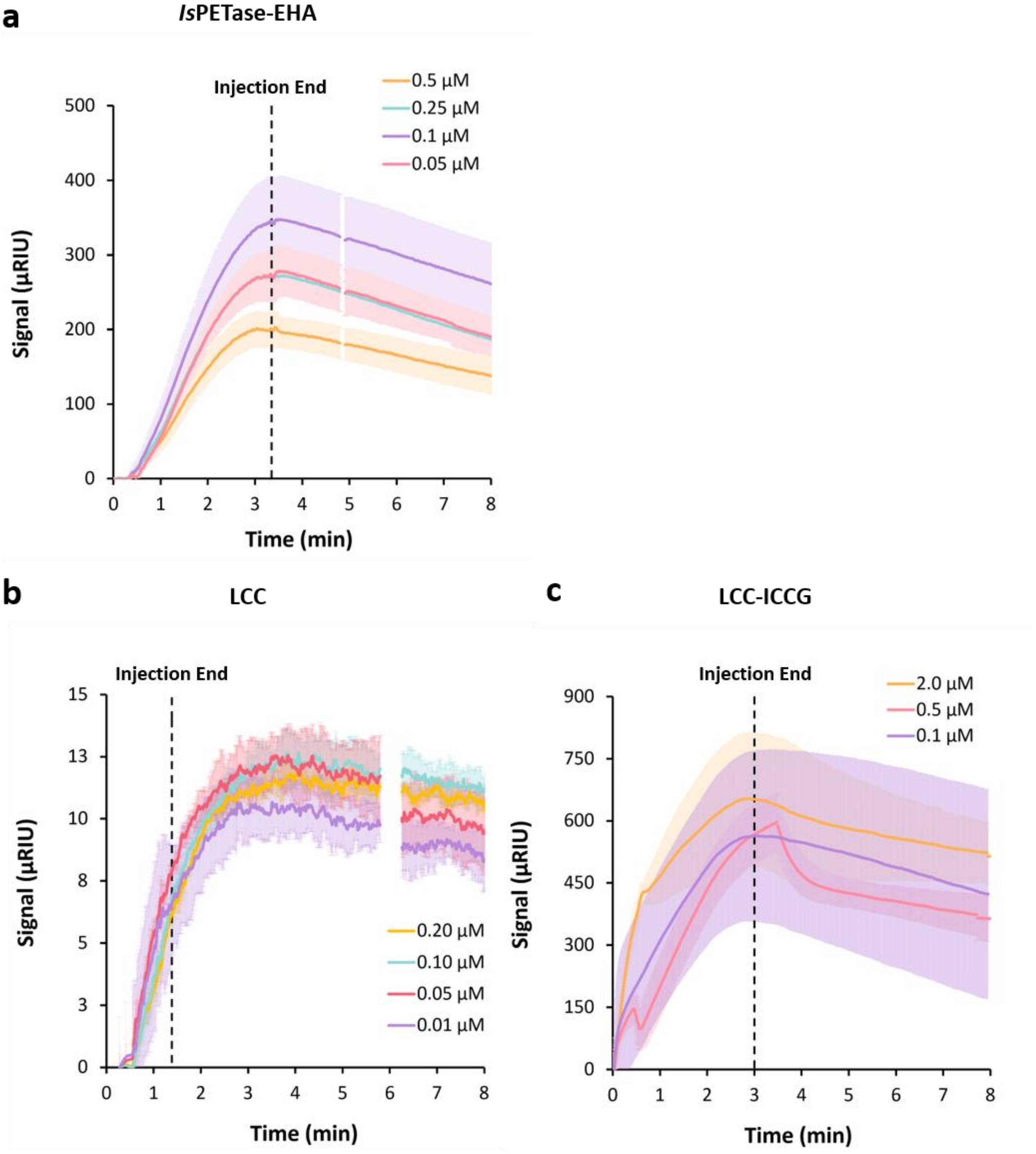


**Supplementary Figure S1. SPR sensorgrams for 3PET binding to immobilised PET hydrolases.** 0.5 µM His-tagged (**a**) IsPETase-EHA, (**b**) LCC, or (**c**) LCC-ICCG was captured on a Ni-NTA SPR chip, and 3PET was injected in 20 mM HEPES pH 8 containing 0.01% (a, b) or 0.05% (c) Triton X-100. Conditions: (a) 0.05–0.5 µM, 25 µL/min, 40 °C, 3 min injection; (b) 0.01–0.20 µM, 45 µL/min, 50 °C, 50 s injection; (c) 0.1–2.0 µM, 25 µL/min, 60 °C, 3 min injection. Dashed lines indicate injection end. Curves are control-channel and buffer-corrected and shading indicates standard deviation across n = 3 injections. Peak response decreases with increasing 3PET concentration in all three cases, inconsistent with a simple binding isotherm and indicative of analog aggregation. These data were therefore not analysed kinetically.

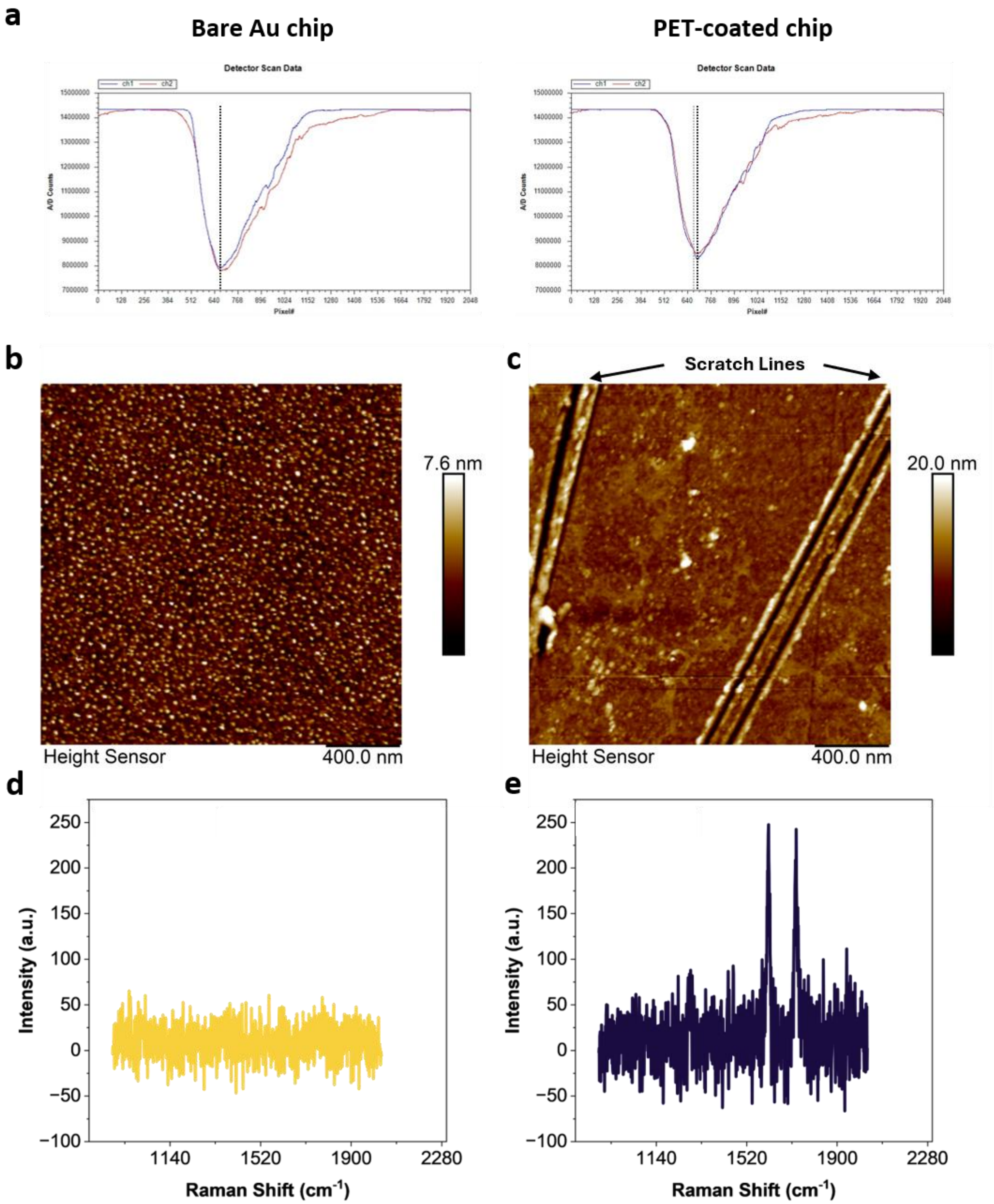


**Supplementary Figure S2. Characterisation of PET-coated SPR chips.** (**a**) SPR detector scans of a bare gold chip (left) and a PET-coated chip (right), acquired in 20 mM HEPES, pH 8 at 25 °C. Both flow channels are shown (ch1, blue; ch2, red). Dashed lines mark the SPR trough minimum. AFM height images of (**b**) a bare gold chip and (**c**) a PET-coated chip, showing loss of the granular gold texture. Diagonal lines in (c) are scratch-test lines, from which a soft-layer thickness of roughly 7 nm was determined. Confocal Raman spectra of (**d**) a bare gold chip and (**e**) a PET-coated chip. Peaks at 1615 and 1730 $cm^{-1}$, assigned to the aromatic ring stretch and the ester carbonyl stretch of PET respectively, are present only on the coated surface.

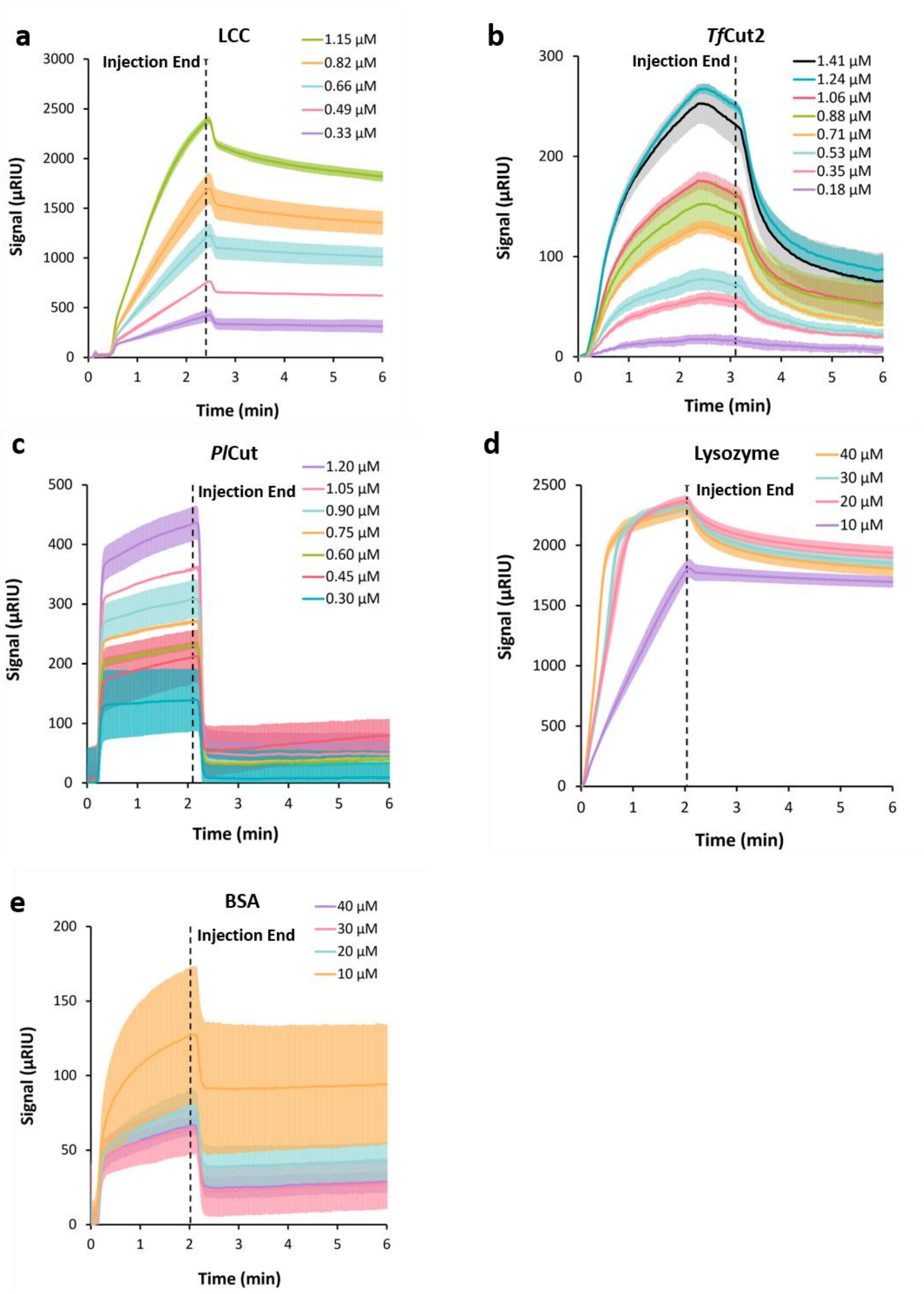


**Supplementary Figure S3. Concentration-dependent SPR sensorgrams for PET hydrolases and non-degrading control proteins on PET-coated chips.** Sensorgrams for (**a**) LCC, (**b**) *Tf*Cut2, (**c**) PlCut, (**d**) lysozyme and (**e**) BSA, injected over PET-coated SPR chips in 20 mM HEPES, pH 8 at 25 µL/min. Enzyme concentrations are given in each legend. Dashed lines indicate the end of the injection. Temperatures were (**a, b**) 50 °C, and (**c** - **e**) 25 °C. Curves are control-channel and buffer-corrected. Shading indicates standard deviation across n = 3 injections.

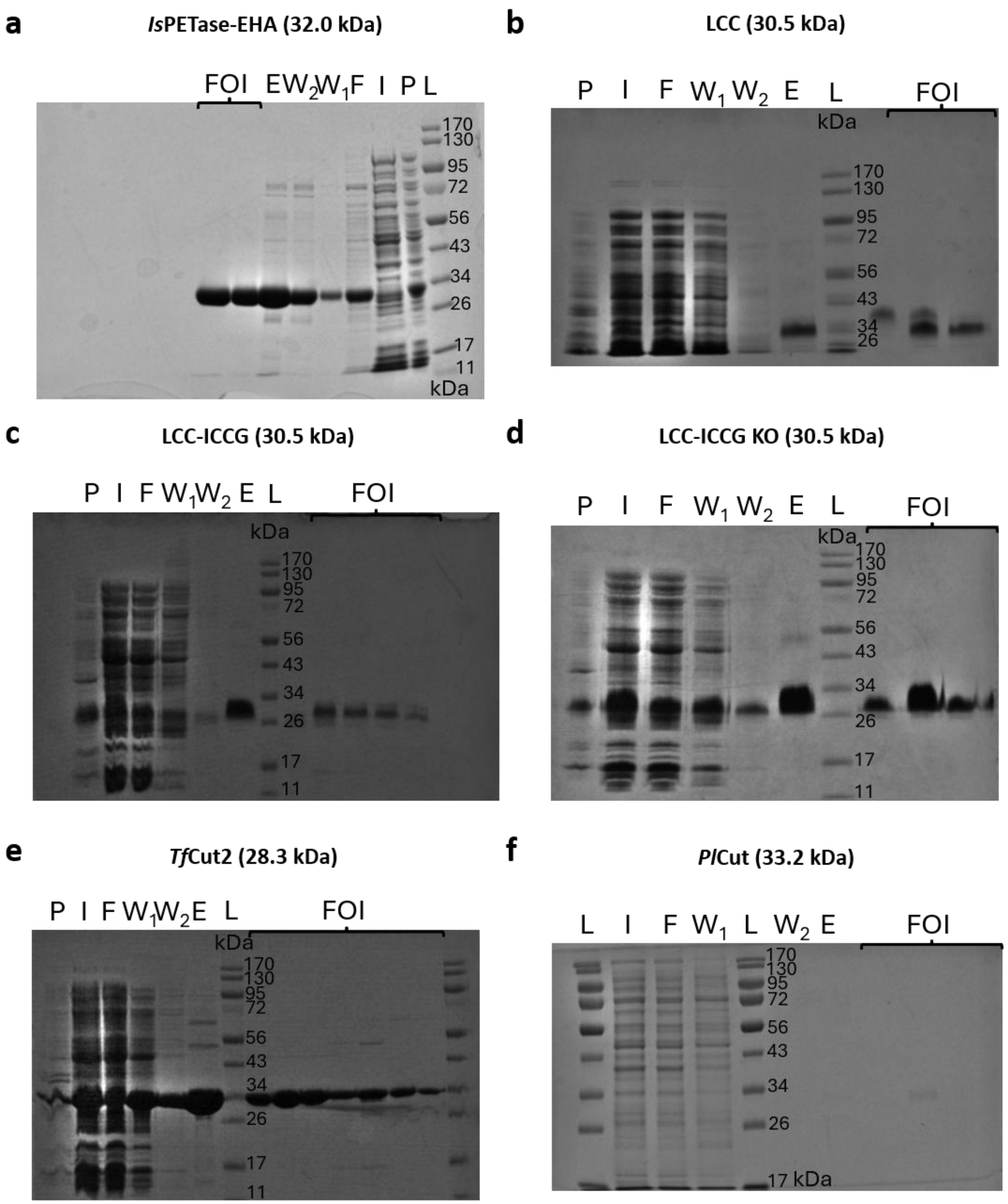


**Supplementary Figure S4. SDS-PAGE analysis of PET hydrolase expression and purification.** Coomassie-stained 12% polyacrylamide gels showing purification of (**a**) *Is*PETase-EHA (32.0 kDa), (**b**) LCC (30.5 kDa), (**c**) LCC-ICCG (30.5 kDa), (**d**) LCC-ICCG S165A (30.5 kDa), (**e**) *Tf*Cut2 (28.3 kDa) and (**f**) *Pl*Cut (33.2 kDa). Lanes: P, insoluble pellet; I, clarified input; F, column flow-through; $W_1$ and $W_2$, wash fractions (100 mM sodium phosphate buffer pH 7.4, 150 mM NaCl, 60 mM imidazole); E, elution (100 mM sodium phosphate buffer pH 7.4, 150 mM NaCl, 300 mM imidazole); L, molecular weight ladder (PageRuler Plus); FOI, pooled size-exclusion fractions of interest. Each enzyme eluted as a single dominant band at the expected molecular weight.

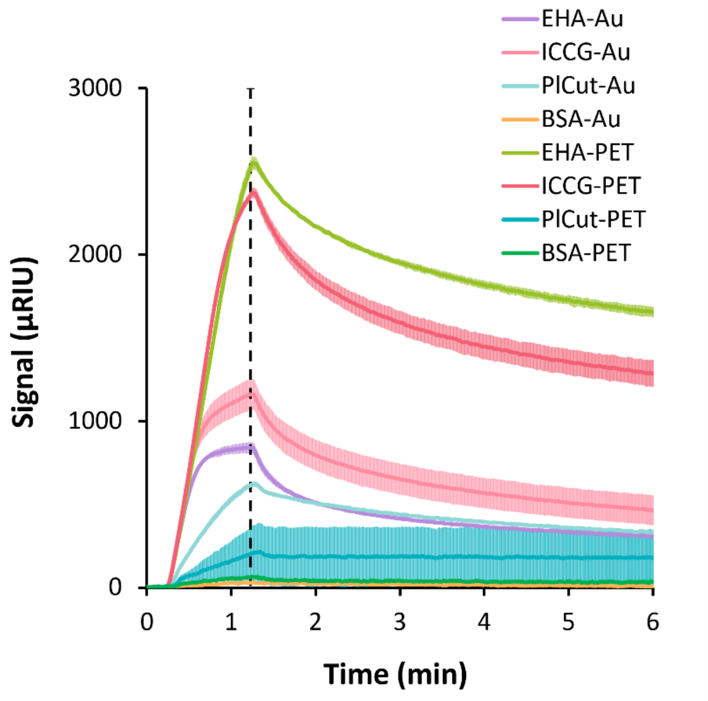


**Supplementary Figure S5. PET hydrolase binding is specific to the PET layer rather than the underlying gold.** 1.0 µM IsPETase-EHA, LCC-ICCG, PlCut and BSA were injected over PET-coated (dark traces) or bare gold (light traces) SPR chips in 20 mM HEPES, pH 8 at 45 °C and 25 µL/min. The dashed line indicates the end of injection. Shading indicates standard deviation across n = 3 injections. Both PET hydrolases gave substantially greater responses on PET than on gold (3-fold for *Is*PETase-EHA, 2-fold for LCC-ICCG), with correspondingly slower dissociation. *Pl*Cut showed the opposite behaviour, binding bare gold approximately 2.6-fold more than PET, indicating that its interaction with hydrophobic surfaces does not extend to productive engagement with the polymer. BSA responses were substantially lower on both surfaces.

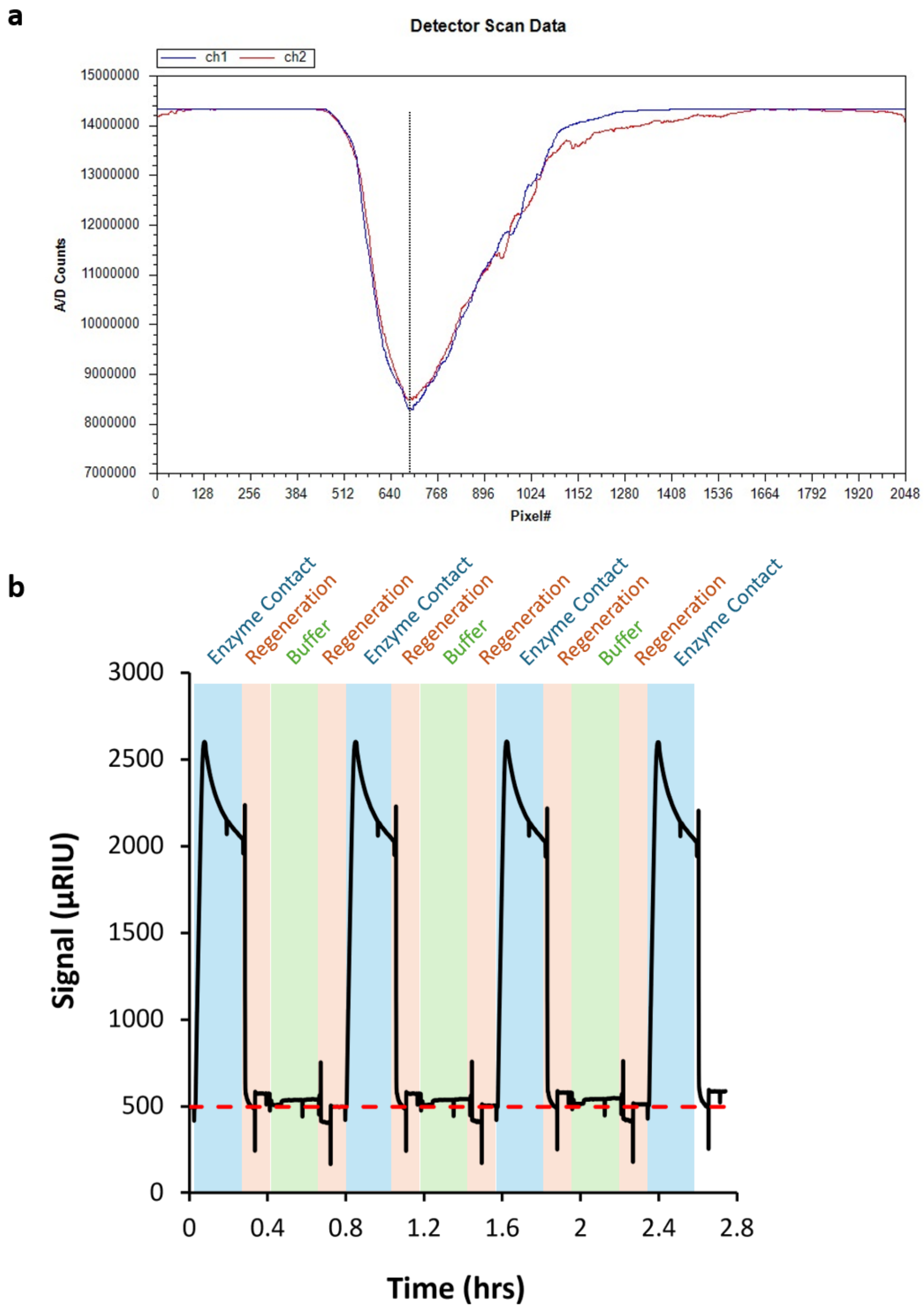


**Supplementary Figure S6. Surface stability across repeated binding and regeneration cycles.** (**a**) SPR detector scan of a PET-coated chip following an injection, showing both flow channels. Channel 1 (blue) was exposed to enzyme throughout the experiment, while channel 2 (red) received buffer only. The dashed line marks the resonance minimum for both channels, indicating no detectable loss of the PET layer. (**b**) Continuous trace of four consecutive cycles of 1.31 µM LCC-ICCG injection, regeneration with 0.1 M NaOH, and buffer equilibration in 20 mM HEPES, pH 8 at 50 °C and 25 µL/min. Shaded regions denote enzyme contact (blue), regeneration (orange) and buffer (green). The dashed red line marks the mean post-regeneration baseline. Baseline and maximum response were reproducible across cycles, indicating that neither enzyme binding nor regeneration measurably alters the PET surface post-dissociation.

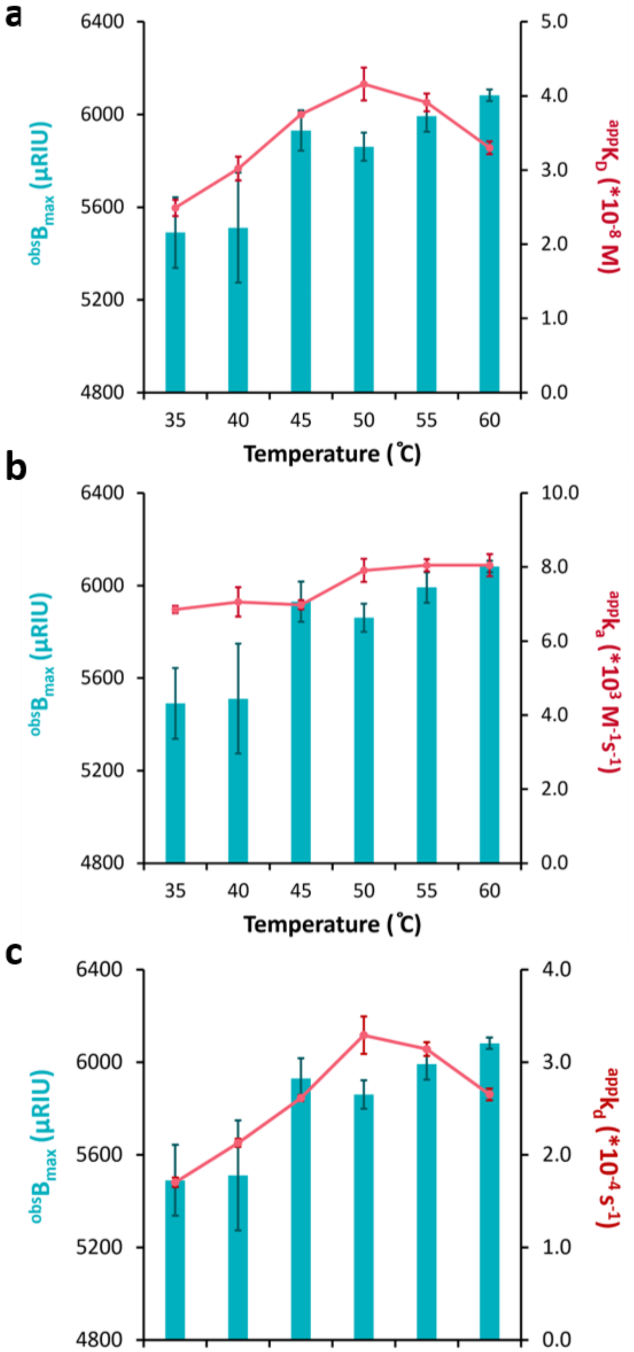


**Supplementary Figure S7. Temperature dependence of IsPETase-EHA binding to a PET-coated surface.** Apparent maximum response ($B_{max}$, orange bars, left axis) shown against (**a**) apparent equilibrium dissociation constant ($K_D$), (**b**) apparent association rate constant ($k_a$), and (**c**) apparent dissociation rate constant ($k_d$) (pink lines, right axes) for 1.0 µM IsPETase-EHA injected at 25 µL/min in 20 mM HEPES pH 8 at 35–60 °C (n = 4 ± 1 SD). $B_{max}$ and $k_a$ increase gradually and monotonically across the range (10.8% and 17.6% respectively), consistent with modestly enhanced surface accessibility or encounter efficiency. In contrast, $k_d$ is non-monotonic, rising to a maximum at 50 °C before declining at 60 °C. This is the primary contributing factor to the temperature dependence of $K_D$, which resembles the non-monotonic $k_d$ relationship. Complex lifetime is therefore governed by the microscopic properties of the bound state rather than by substrate accessibility alone.

**a**

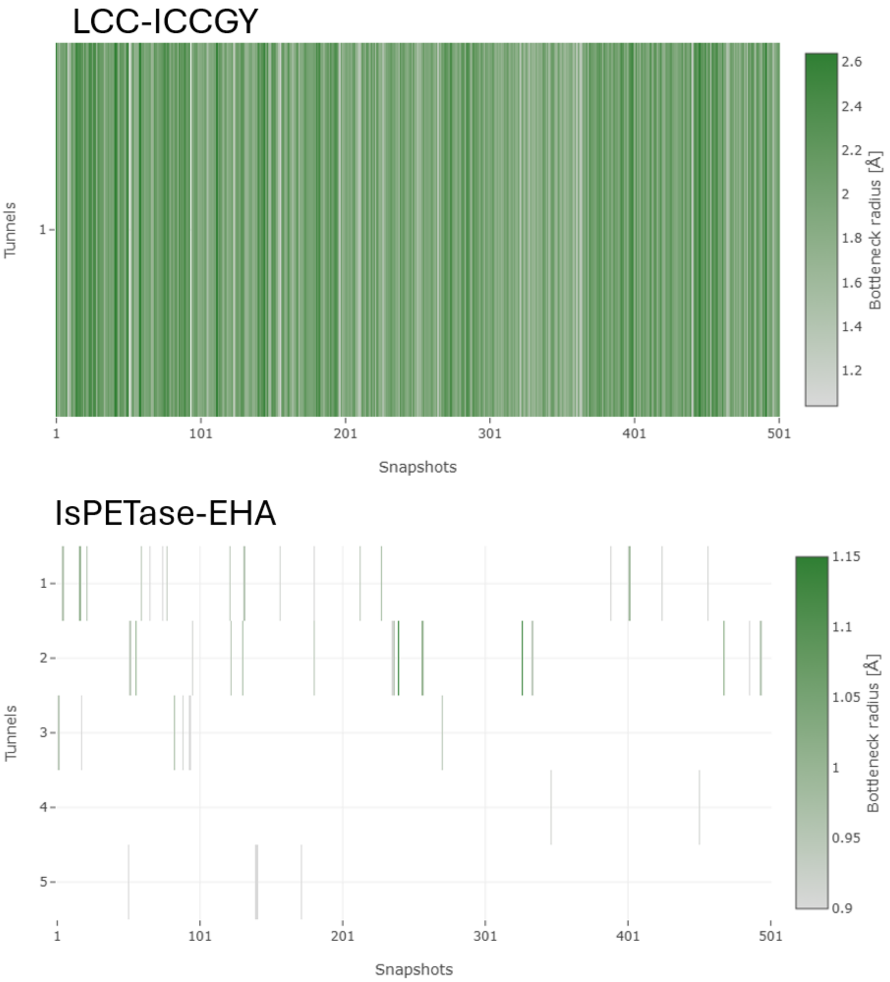


**c**

| Enzyme | Occupancy | Bottleneck radius (Å) | Maximum bottleneck radius (Å) | Length (Å) | Curvature | Throughput |
|---|---|---|---|---|---|---|
| LCC-ICCGY | 499 | 1.99 ±0.27 | 2.64 | 1.99 ±0.86 | 1.08 ±0.11 | 0.91 ±0.02 |
| IsPETase-EHA | 17 | 0.94 ±0.03 | 1 | 10.97 ±1.57 | 1.34 ±0.16 | 0.45 ±0.04 |
| | 15 | 0.98 ±0.07 | 1.15 | 20.2 ±2.36 | 1.23 ±0.07 | 0.28 ±0.05 |
| | 6 | 0.93 ±0.02 | 0.97 | 18.04 ±1.63 | 1.38 ±0.09 | 0.31 ±0.04 |
| | 2 | 0.91 ±0 | 0.91 | 14.3 ±0 | 1.3 ±0.02 | 0.34 ±0 |
| | 4 | 0.9 ±0 | 0.9 | 36.62 ±2.43 | 1.63 ±0.05 | 0.07 ±0.01 |

**Supplementary Figure S8. Transient channel formation in LCC-ICCGY and IsPETase-EHA.** Channels were identified using CAVER across 501 snapshots of (**a**) LCC-ICCGY (PDB 8QRJ) and (**b**) IsPETase-EHA (PDB 6IJ6), using a probe radius of 0.9 Å and a starting point defined by residues 97, 159, 182, 183, 201–203, 230, 232, 237–241, and 249–250 for 6IJ6 and 93–97, 101–102, 164–165, 212, 242–243, and 246 for 8QRJ. Each vertical line indicates a snapshot in which the corresponding channel was detected, shaded by bottleneck radius. (**c**) Channel parameters are reported as mean ± 1 SD across the snapshots in which each channel was present. LCC-ICCGY presents a single channel detected in 499 of 501 snapshots, forming part of the substrate-binding cleft. The more open cleft of IsPETase-EHA does not meet CAVER's channel detection criteria and is therefore not represented. The channels identified in (b) are instead transient openings elsewhere in the structure, each present in fewer than 4% of conformations and with bottleneck radii near the detection limit. The greater number of such transient channels in IsPETase-EHA is consistent with the increased conformational sampling inferred from its binding thermodynamics and surface kinetics.

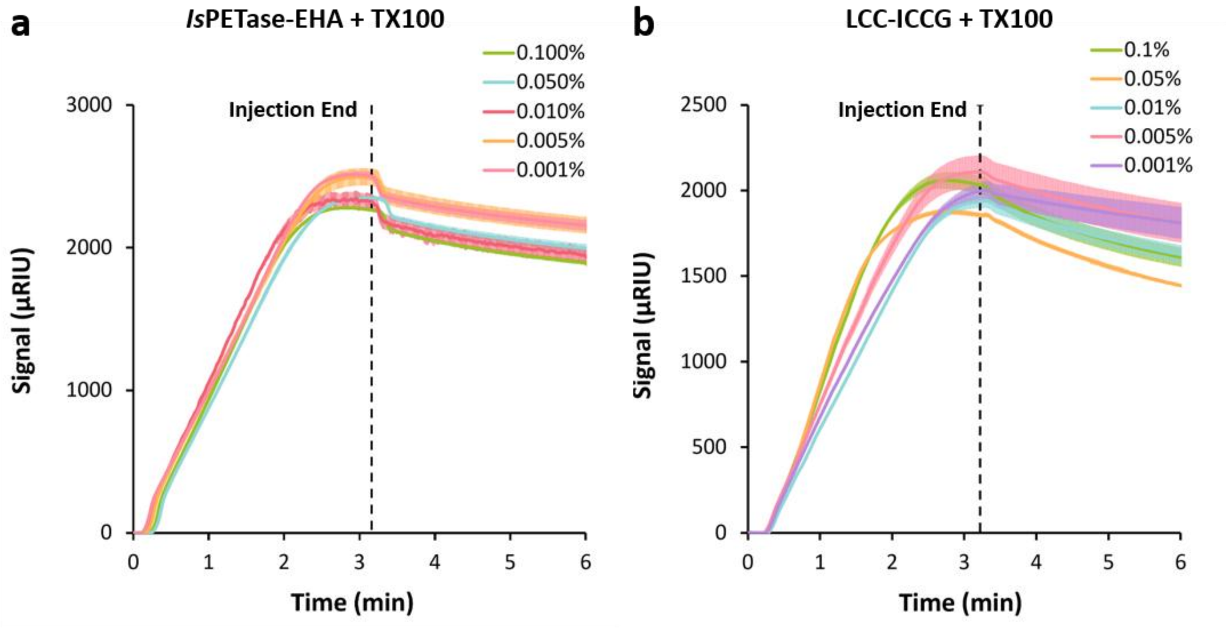


**Supplementary Figure S9. Effect of stabilizers on SPR sensorgrams for PET hydrolases.** Sensorgrams for 1.0 µM (**a**) *Is*PETase-EHA, and (**b**) LCC-ICCG injected over PET-coated SPR chips in 20 mM HEPES, pH 8 at 25 µL/min with added TX100 at concentrations from 0.001 to 0.1 %v/v. Dashed lines indicate the end of the injection. Temperatures were 45 °C (*Is*PETase-EHA), or 60 °C (LCC-ICCG). Curves are control-channel and buffer-corrected. Shading indicates standard deviation across n = 3 injections.

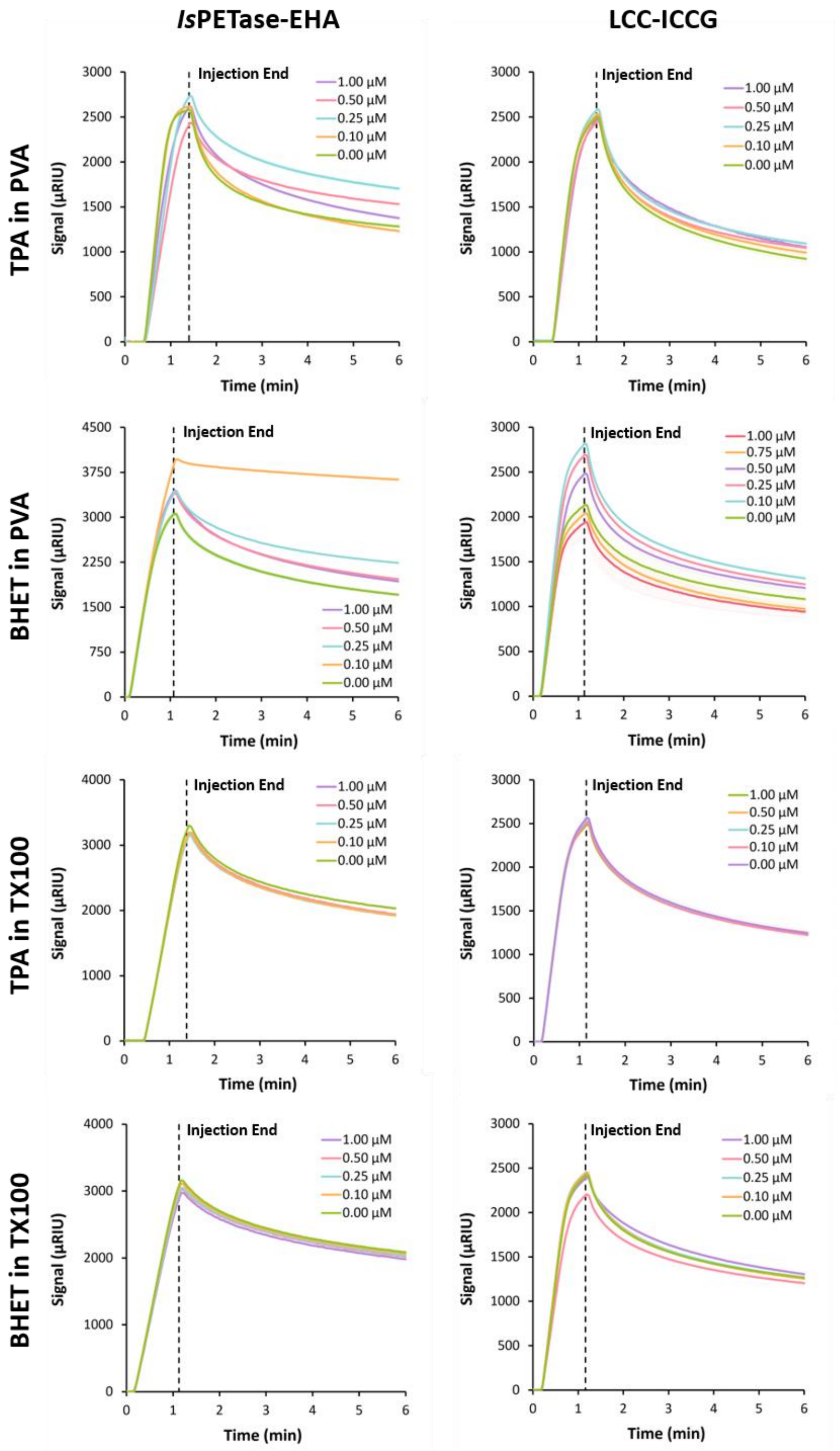


**Supplementary Figure S10. Degradation products modulate PET hydrolase surface engagement in PVA but not in Triton X-100.** Sensorgrams for 1.0 µM *Is*PETase-EHA (left) and LCC-ICCG (right) injected over PET-coated chips in 20 mM HEPES pH 8 containing 0.01% PVA (rows 1, 2) or 0.01% Triton X-100 (rows 3, 4), in the presence of 0 – 1.0 µM TPA (rows 1, 3) or BHET (rows 2, 4), at 45 °C (*Is*PETase-EHA) and 60 °C (LCC-ICCG), at 25 µL/min. Dashed lines indicate the end of injection. Shading indicates standard deviation across n = 3 injections. In PVA, both products alter surface engagement biphasically, with progressive loss of the effect at higher concentrations. In Triton X-100, the curves are near-superimposable across the full product range for both enzymes and both products, indicating that this surfactant suppresses the product effect.
